# A promiscuous archaeal cardiolipin synthase generating a variety of cardiolipins and phospholipids

**DOI:** 10.1101/2020.12.02.408559

**Authors:** Marten Exterkate, Niels A. W. de Kok, Ruben L. H. Andringa, Niels H. J. Wolbert, Adriaan J. Minnaard, Arnold J. M. Driessen

**Author notes:** To whom correspondence should be addressed: Arnold J. M. Driessen: Department of Molecular Microbiology, Groningen Biomolecular Sciences and Biotechnology Institute and Zernike Institute for Advanced Materials, University of Groningen, Nijenborgh 7, 9747 AG, Groningen, The Netherlands.

## Abstract

Cardiolipin (DPCL) biosynthesis has barely been explored in Archaeal isoprenoid-based ether lipid membranes. Here, we identified a cardiolipin synthase (MhCls) from the mesophilic anaerobic methanogen *Methanospirillum hungatei.* The enzyme was overexpressed in *Escherichia coli,* purified, and subsequently characterized by LC-MS. MhCls utilizes two archaetidylglycerol molecules in a transesterification reaction to synthesize archaeal di-phosphate cardiolipin (aDPCL) and glycerol. The enzyme is invariant to the stereochemistry of the glycerol-backbone and the nature of the lipid tail, as it also accepts phosphatidylglycerol to generate di-phosphate cardiolipin (DPCL). Remarkably, in the presence of archaetidylglycerol and phosphatidylglycerol, MhCls formed an archaeal-bacterial hybrid di-phosphate cardiolipin (hDPCL), that so far has not been observed in nature. Due to the reversibility of the transesterification, cardiolipin can be converted back in presence of glycerol into phosphatidylglycerol. In the presence of other compounds that contain primary hydroxyl groups (e.g. alcohols, water, sugars) various natural and unique artificial phospholipid species could be synthesized, including multiple di-phosphate cardiolipin species. Moreover, MhCls could utilize a glycolipid in the presence of phosphatidylglycerol to form a glycosyl-mono-phosphate cardiolipin, emphasizing the promiscuity of this cardiolipin synthase.

## Introduction

Cardiolipin (CL), or more specifically di-phosphate cardiolipin (DPCL), is present in lipid membranes throughout all three domains in life. DPCL is usually a relatively small component (< 10 mol%) of the total membrane lipid composition, and its primary role appears to be supporting the function of various membrane proteins (1, 2). Unlike most naturally occurring glycerophospholipids, DPCL consists of two 1,2-diacylphosphatidate moieties esterified to the 1- and 3-hydroxyl groups of a glycerol molecule. Because of the two phosphates it can carry up to two negative charges (3). Furthermore, the polar head group is relatively small compared to the four bulky hydrophobic tails, giving cardiolipin its characteristic inverted conical shape in the presence of divalent cations (4). Due to this structural feature, DPCL may induce membrane curvature. Indeed, DPCL is believed to be located in lipid-domains at the cell poles and division site in bacteria (5–8), and in eukaryotes, it is an important constituent of the curvy mitochondrial membrane (9–11). Besides a tight association with cytochrome *c* oxidase, a part of the respiratory chain complex (12–14), other specific cardiolipin-protein interactions seem not to be conserved among the three domains of life. In bacteria, cardiolipin is often not an essential membrane constituent, as in several organisms it is only produced under certain specific circumstances, such as during stationary phase, or when cells are exposed to certain stressors, e.g. osmotic shock (15–18).

Cardiolipin has also been described in archaea, most notably in Euryarchaeota. Specifically, in halophiles the production of archaeal di-phosphate cardiolipin (aDPCL) is influenced by changes in the ionic composition of the environment (19). For instance, in the halophilic organism *Halorubrum sp.*, hypotonic stress resulted in increased production of aDPCL, as well as the glycocardiolipin S-di-glycosyl-archaeal mono-phosphate cardiolipin (S-2glyco-aMPCL, originally referred to as: S-DGD-5-PA), which consists of an archaetidic acid (AA) molecule attached to a sulfated diglycosyl diphytanylglycerol diether (S-2Glyco-DGD) (20). Further, a variety of other glycosyl-mono-phosphate cardiolipin (glyco-MPCL) species have been identified in archaea (19). For example, *Halobacterium salinarum* produces a S-tri-glycosyl-diether glycolipid fused to AA (S-3Glyco-aMPCL, originally referred to as: S-TGD-1-PA) (21), while *Haloferax volcanii* also contains the glycosyl cardiolipin analogue S-2Glyco-aMPCL (originally referred to as: S-GL-2) (22). The polar head group of these glyco-MPCL species is structurally very different from the classical glycerol-based di-phosphate DPCL, which raises questions on the specific function of these cardiolipin species, as well as the enzymatic mechanism of their synthesis.

Whereas cardiolipin synthesis in eukaryotes and bacteria has been studied extensively, the mechanism of cardiolipin biosynthesis in archaea has largely remained unexplored. In eukaryotes, di-phosphate cardiolipins are synthesized from the substrates cytidine diphosphate diacylglycerol (CDP-DAG) and phosphatidylglycerol (PG) (23). In this reaction, the first phosphate group connected to DAG is coupled to the polar glycerol head of PG, and a cytidine monophosphate (CMP) is released. On the other hand, in bacteria DPCL is most commonly synthesized by transferring the phosphatidyl-group, originating from a PG molecule, to another PG, whereby a glycerol molecule is released (24). This reaction is catalyzed by the ClsA and ClsB-type cardiolipin synthases (Cls) (25, 26). However, an alternative synthesis method has been identified in *Escherichia coli* in which phosphatidylethanolamine (PE) is used together with PG, to form cardiolipin and ethanolamine, a reaction catalyzed by the ClsC-type enzymes (17). Cardiolipin synthases belong to a superfamily of membrane proteins that harbor phospholipase D motifs. Homology searches in genomes of archaea revealed Cls-like members that belong to the CL superfamily. However, until now no archaeal enzyme has been identified or experimentally associated with cardiolipin biosynthesis. Here, we report on the identification and characterization of a cardiolipin synthase of the Euryarchaeote *Methanospirillum hungatei*. The enzyme showed a remarkable promiscuity with respect to accepted and produced lipid species, ranging from a wide variety of phospholipids and di-phosphate cardiolipins to include even glycolipid and glycosyl-mono-phosphate cardiolipin species.

## Results

### Bioinformatic identification of cardiolipin synthases in Archaea

Cardiolipin-like molecules have only been identified in some archaea, most notably in halophiles; a group of organisms belonging to the phylum of the Euryarchaeota (19). To identify possible cardiolipin synthesizing enzymes in archaea, we performed a BLAST homology search to each of the three Cls enzymes (ClsA, ClsB, and ClsC) identified in the bacterium *Escherichia coli* as template. This resulted in similar hits, with the best sequence coverage and identity with ClsA (Figure S1). Although the structure of this membrane protein has not been determined, a membrane topology prediction based on the average hydropathy-profile of the amino acid sequences of a family of bacterial ClsA proteins reveals the presence of two hydrophobic regions at the N-terminus (corresponding to amino acids 7-29 and 39-61 in ClsA) (Figure 1A). These could represent transmembrane segments that are linked to a C-terminal globular domain (Figure 1B, left panel) that contains two HKD motifs (HxKx_4_D), which are universally present in cardiolipin synthesizing enzymes, and a common characteristic for enzymes of the phospholipase D superfamily (26). The BLAST homology search revealed 2 main clusters of archaeal homologs that also contain the two HKD motifs (Figure 1C). One cluster consists of halophilic archaea and the other concerns methanogenic archaea, both groups belonging to the Euryarchaeota. The enzymes from the cluster of methanogenic archaea show about 25% sequence identity with the *E. coli* ClsA, which include the predicted two hydrophobic regions at the N-terminus (Figure S2). Additionally, the second predicted N-terminal hydrophobic domain show conserved residues forming the motif: Wx_7_Px_2_Gx_3_Yx_3_G (“x” represents a hydrophobic amino acid), a feature which is also conserved among bacterial ClsA-type proteins (Figure 1D). The high incidence of hydrophobic amino acids suggests that this region is likely embedded in the membrane. The conserved proline and glycine (PxxG) residues are located in the middle of the hydrophobic stretch. These two amino acids are often found in turns and loops and thus may introduce flexibility in this predicted helix-region (27, 28). Furthermore, this hydrophobic region is flanked on both sides by multiple positive charges, which are known to inhibit translocation across the membrane according to the positive-inside rule, and thus may affect the membrane topology of this region. As a consequence, this predicted helix-domain may not span across the membrane, but represent a re-entrance loop that enters and leaves the membrane at the same leaflet side (Figure 1B, right panel). Taken together, this group of methanogenic ClsA-like proteins appear to be promising candidates for potential archaeal cardiolipin synthases. The putative *Methanospirillum hungatei* Cls (WP_011448254) was selected for further characterization, as it grows at mesophilic temperatures and a pH around 7, with an osmotic requirement comparable to that of *E. coli*.

**Figure 1.**
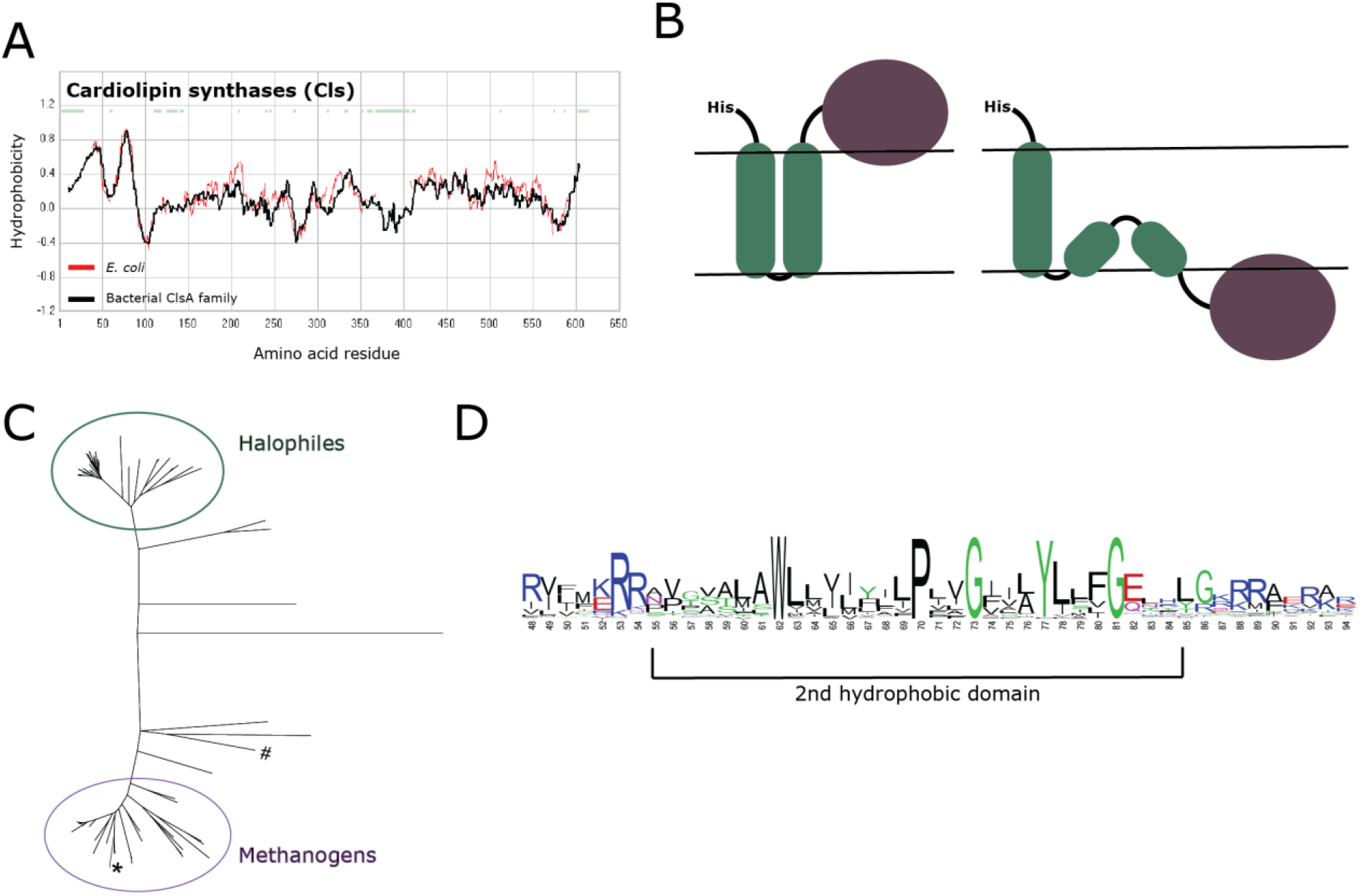
Bioinformatic identification of an archaeal cardiolipin synthase (Cls). **(a)** Hydropathy profile alignment of *E. coli* ClsA (red line), with the averaged hydropathy profile of its bacterial protein family (black line). **(b)** Schematic representation of potential membrane topologies of cardiolipin synthase with 2 predicted transmembrane anchors and a globular active domain. **(c)** Unrooted tree of putative archaeal cardiolipin synthases with 2 main clusters. # *E. coli* ClsA; * *M. hungatei* Cls. **(d)** Consensus sequence logo of the second hydrophobic region of cardiolipin synthase type A enzymes from bacteria and the group of methanogenic archaea.

### Archaeal cardiolipin synthesis from archaetidylglycerol

The putative *M. hungatei* ClsA homologue (MhCls) was ordered as an *E. coli* codon-optimized synthetic gene, cloned into a His-tag containing overexpression vector and expressed in *E. coli*. Overexpressed MhCls was recovered from the membrane fraction, solubilized with the detergent n-dodecyl-β-d-maltoside (DDM; 2%, w/v), and purified by Ni-NTA agarose affinity chromatography (Figure 2A). To confirm that MhCls is indeed an archaeal cardiolipin synthase, the activity of purified enzyme towards the substrate archaetidyl glycerol (AG) was tested *in vitro*. Since this archaeal equivalent of phosphatidylglycerol (PG) is not commercially available, both the *sn*1-*sn*1 **(14)** and the *sn*1-*sn*3 **(14’)** diastereomers of AG were chemically synthesized (Figure 2B) (see the supplemental information). The bis-phytanyl glycerol core was readily synthesized from commercially available phytol and glycidol. Both enantiomers of the glycerol-phosphate headgroup were also produced by asymmetric synthesis and were individually coupled to the aforementioned chiral lipid core via a phosphor-amidite coupling. After the successful synthesis of *sn1-sn1* and *sn1-sn3* AG, first a mixture of both diastereomers was incubated overnight at 37°C together with purified MhCls. LC-MS analysis revealed that the majority of AG was consumed which coincided with the production of an ion *m/z* 1520.30 [M-H]^−^, corresponding to archaeal di-phosphate cardiolipin (aDPCL) (Figure 2C; for fragmentation data see Figure S4). In the absence of MhCls no such conversion was observed. Subsequently the activity of MhCls towards the individual AG diastereomers (*sn1* and *sn3*) was tested and compared, but both acted as substrates in a similar manner (data not shown). Aside from producing aDPCL, MhCls additionally synthesized the lipid species archaetidic acid (AA) (Figure 2C). The latter is probably the result of an unsuccessful transfer of the archaetidyl-group, whereby the enzyme hydrolyzes the terminal phosphodiester bond in a phospholipase D-like manner. Overall, it can be concluded that MhCls is an archaeal cardiolipin synthase.

**Figure 2.**
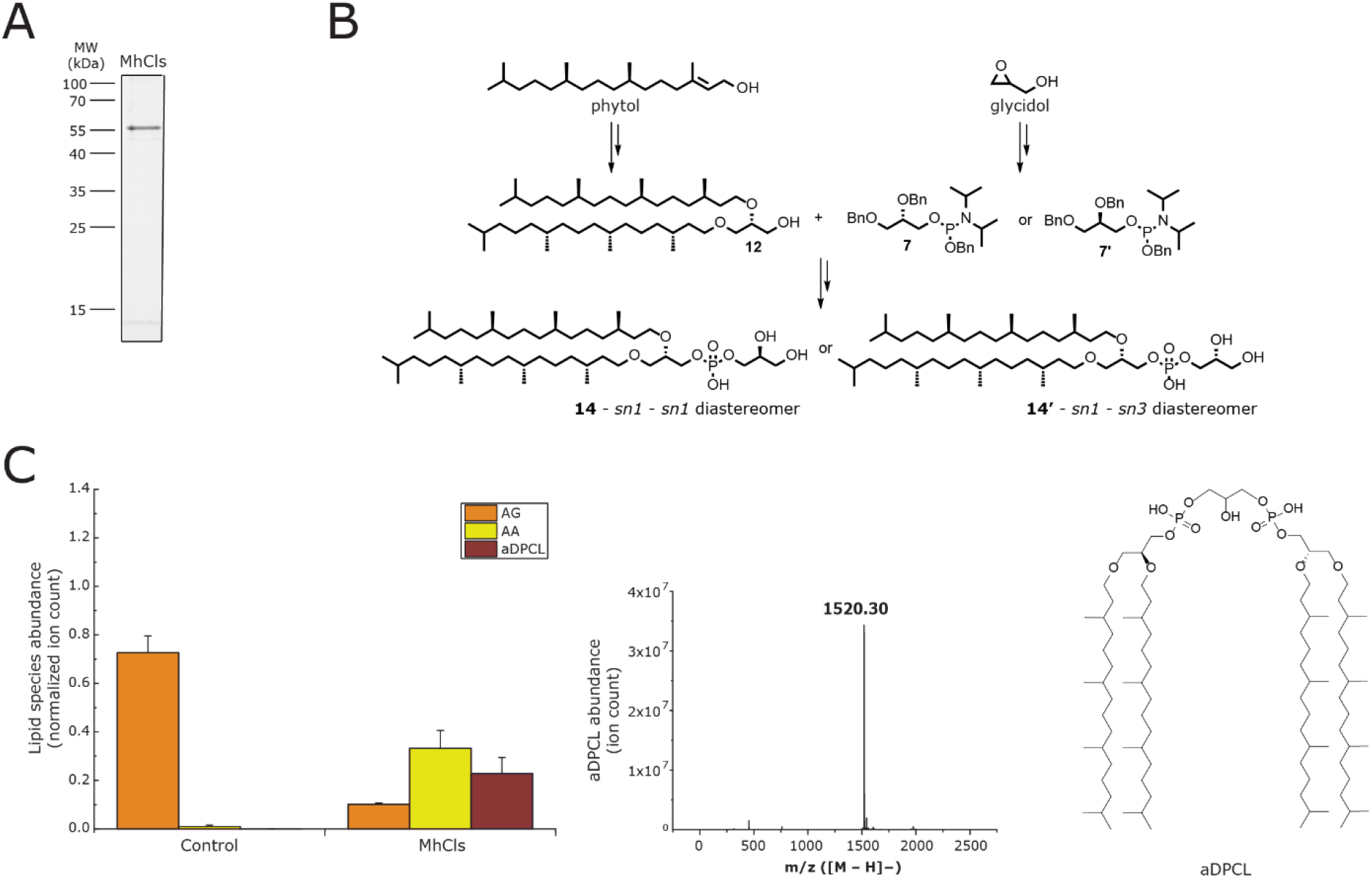
Purification and activity of a cardiolipin synthase from *M. hungatei* (MhCls). **(a)** Coomassie stained SDS-PAGE gel of the MhCls purified by Ni-NTA chromatography. (b) Schematic representation of chemical AG synthesis. **(c)**. *In vitro* activity of purified MhCls. Lipid species were analyzed by LC-MS, normalized for the internal standard DDM, and plotted on the y-axis. Mass spectrum showing the presence of archaeal cardiolipin (aDPCL) as *m/z* 1520.30 [M-H]^−^), and its structure.

### A glycerol-dependent dynamic equilibrium between PG and DPCL

Next, the activity of MhCls towards the bacterial equivalents of these lipids was tested. To perform the measurements under optimal conditions, adequate membrane reconstitution of MhCls is required for which a bacterial lipid/detergent molar ratio-profile was recorded for the substrate PG (Figure S5). Subsequently, MhCls activity was monitored in the presence of PG, which resulted in the formation of DPCL, showing that the enzyme accepts both bacterial and archaeal phospholipids as a substrate. Similar to AG, utilization of PG not only resulted in the production of DPCL, but also PA. Initially most of the PG is converted into DPCL, although some PA is produced as well (Figure 3A, purple lines). Eventually the PG levels reach a plateau, and DPCL levels start to drop, concomitantly with the continuous production of PA. In the presence of a high concentration (100 mM) of glycerol (Figure 3A, green lines), the initial conversion of PG into DPCL is similar, but production of PA is significantly reduced, while PG reaches a plateau level at a higher concentration. This indicates that glycerol most likely stimulates the formation of PG in a reverse transesterification reaction, and thereby competes with the hydrolysis of DPCL (Figure 3D). To confirm this hypothesis, the same reaction was performed with DPCL as substrate. In the presence of 100 mM glycerol (Figure 3B, green lines), DPCL is initially predominantly converted into PG, while only low levels of PA are noted. In contrast, in the absence of glycerol, PA formation is stimulated, while lower levels of PG are observed (Figure 3B, purple lines). These data demonstrate that PA is formed together with PG by hydrolysis of DPCL (Figure 3D).

**Figure 3.**
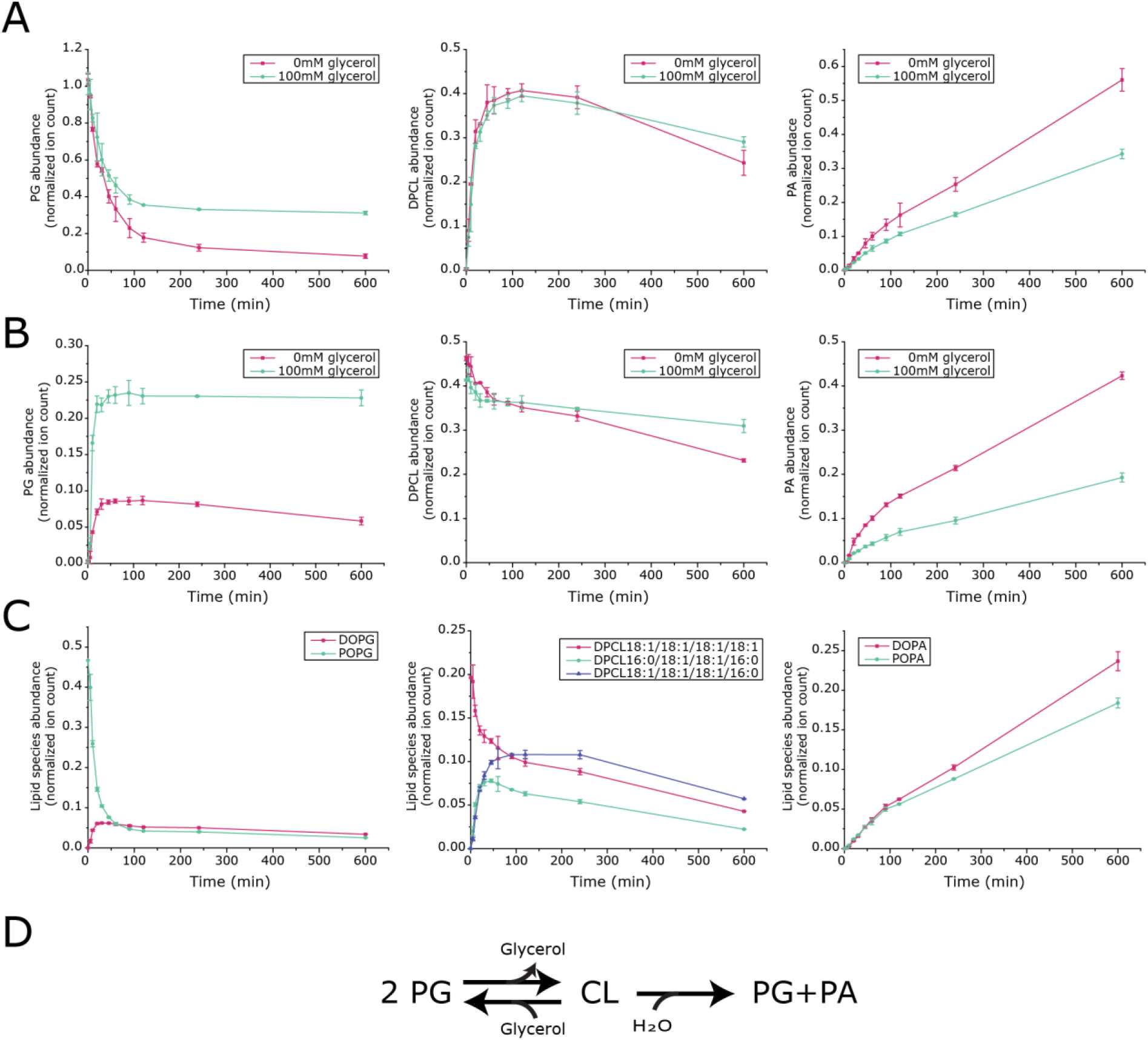
MhCls activity in the presence or absence of glycerol starting with the substrate(s) **(a)** PG, **(b)** DPCL and **(c)** POPG together with DPCL18:1/18:1/18:1/18:1. Lipid species were analyzed by LC-MS, normalized for the internal standard DDM and ion counts are plotted on the y-axis. **(d)** Schematic representation of the ClsA mediated glycerol-dependent dynamic equilibrium in cardiolipin formation.

Hydrolysis of a cardiolipin molecule should yield stoichiometric amounts of PA and PG, but released PG can be directly re-utilized for production of DPCL. To examine the reaction in more detail and to further address the transesterification step, PG and cardiolipin, both with a different acyl-chain composition, were introduced. This enables the detection of a single lipid type as a substrate or a product based on the detected mass. Incubation of MhCls with an equimolar ratio of palmitoyl-oleoyl phosphatidylglycerol (POPG; 16:0/18:1) and di-oleoyl-di-oleoyl di-phosphate cardiolipin (DPCL 18:1/18:1/18:1/18:1), resulted in the expected production of palmitoyl-oleoyl-palmitoyl-oleoyl di-phosphate cardiolipin (DPCL 16:0/18:1/18:1/16:0), di-oleoyl phosphatidylglycerol (DOPG; 18:1/18:1) and di-oleoyl phosphatidic acid (DOPA; 18:1/18:1), but also in the production of palmitoyl-oleoyl phosphatidic acid (POPA; 16:0/18:1), demonstrating that the enzymatic reaction occurs in both directions (Figure 3C). Moreover, a cardiolipin species with the mixed 16:0/18:1 and 18:1/18:1 acyl-chain configuration could be identified as well, which is the synthesis product of POPG combined with DOPG. The latter originated from the reverse-transesterification reaction of DPCL 18:1/18:1/18:1/18:1, thereby showing the dynamic character of the PG-CL equilibrium (Figure 3D). On the other hand, the accumulation of PA suggests that this reaction eventually becomes unidirectional as PA cannot be further used. Indeed, when POPA (16:0/18:1) was added to a reaction containing DOPG (18:1/18:1), the MhCls-mediated reaction resulted in the formation of only DPCL 18:1/18:1/18:1/18:1 and DOPA, whereas no POPG or DPCL 16:0/18:1/16:0/18:1 was detected, showing that PA cannot be reutilized by MhCls (Figure S6).

### Formation of a bacterial-archaeal hybrid cardiolipin species

Like most archaeal lipids, AG consists of two isoprenoid chains that are coupled to a glycerol-1-phosphate (G1P) backbone via an ether-bond, whereas PG, present in bacteria and eukaryotes, exists of fatty acid tails that are ester-linked to a glycerol-3-phosphate (G3P). This makes that the glycerol backbone of the archaeal AG has the opposite chirality compared to bacterial/eukaryotic PG. To further examine the lipid specificity of MhCls, its activity was examined in the presence of a mixture containing both PG and AG (Figure 4A). Remarkably, not only DPCL and aDPCL were produced, but an additional cardiolipin species was detected: *m/z* 1488.16 [M-H]^−^, that contains one set of isoprenoid lipid-tails and one set of fatty acid lipid-tails (Figure 4B-C; for fragmentation data see Figure S4). This lipid species represents a unique archaeal-bacterial hybrid di-phosphate cardiolipin (hDPCL). Moreover, hDPCL seems to be the major produced CL-species, while about the same amount of ions are detected for aDPCL and DPCL, suggesting that there is no clear preference for any of the lipid substrates PG or AG.

**Figure 4.**
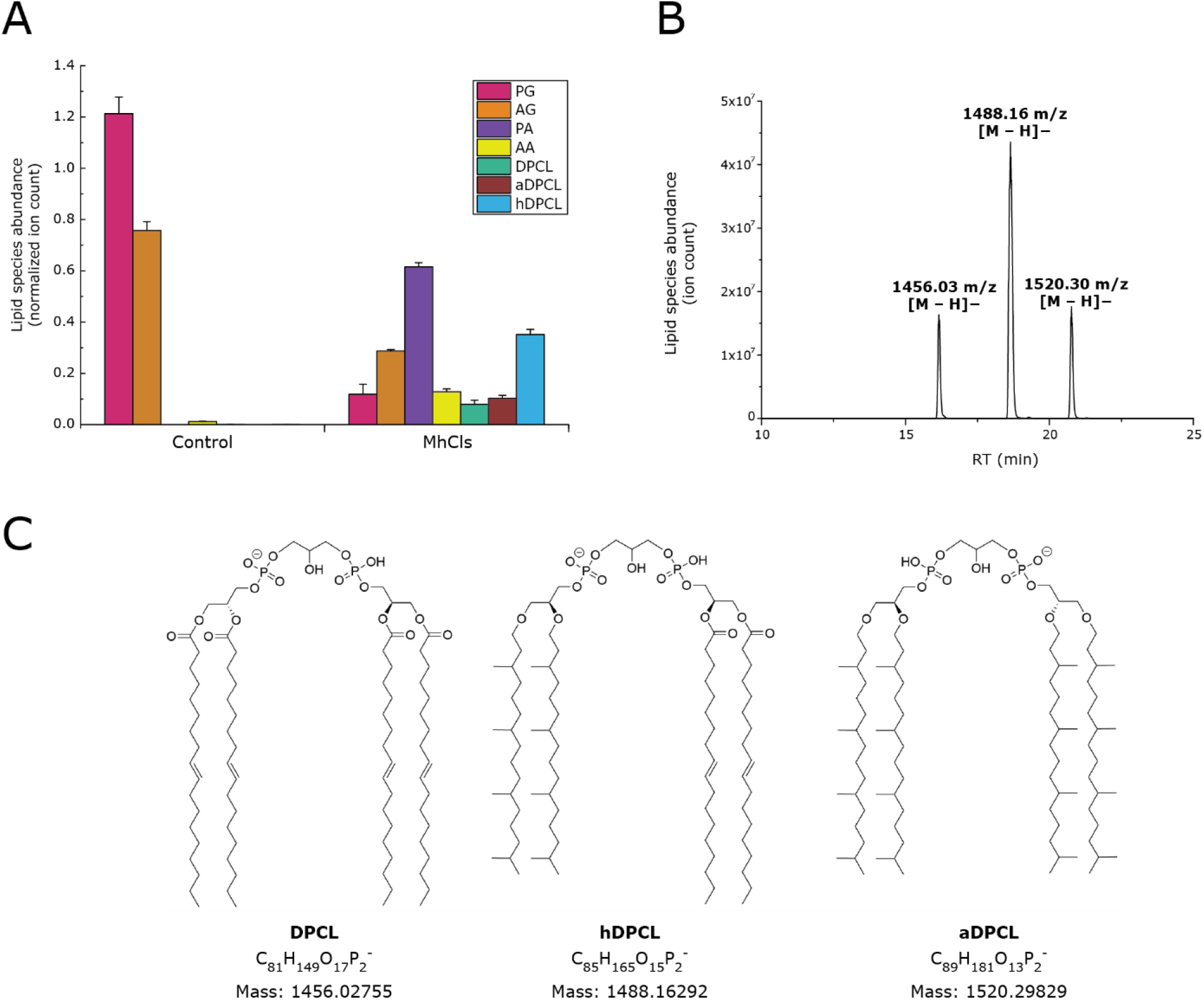
Synthesis of a bacterial-archaeal hybrid cardiolipin species. **(a)** Activity of the bacterial MhCls in the presence of both AG and PG. Lipid species were analyzed by LC-MS, normalized for the internal standard, and plotted. **(b)** LC-MS analysis showing the separation of the produced bacterial cardiolipin (DPCL), hybrid cardiolipin (hDPCL), and archaeal cardiolipin (aDPCL). **(c)** Structures of the three cardiolipin species.

### Diverse polar headgroup incorporation

In the MhCls mediated conversion of DPCL, either glycerol or H_2_O is utilized. This raises the question which other molecules can be used by this enzyme. Therefore, molecules structurally related to glycerol were tested in the DPCL transesterification reaction (Figure 5A). In the presence of 1-propanol, DPCL consumption resulted in the production of substantial amounts of phosphatidyl-1-propanol (P-1-PrOH). On the other hand, the addition of 2-propanol resulted only in low levels (~1%) of phosphatidyl-2-propanol (P-2-PrOH), illustrating the importance of a primary hydroxyl (–OH) group for the transesterification. Similar to 1-propanol, 1,2-propanediol and 1,3-propanediol functioned together with DPCL as substrates for MhCls, resulting in the formation of phosphatidyl-1,2-propanol (P-1,2-PrOH) and phosphatidyl-1,3-propanol (P-1,3-PrOH), respectively. However, in the case of 1,3-PrOH, a clear additional ion *m/z* 1440.20 [M-H]^−^ could be identified, which corresponds to the molecule 1,3-propanediol-di-phosphate cardiolipin (1,3-PrOH-DPCL), a cardiolipin analogue containing a propanediol head group instead of a glycerol. This remarkable cardiolipin-like species could have only been formed if P-1,3-PrOH functioned as a phosphatidyl-acceptor instead of PG in the cardiolipin forming reaction.

**Figure 5.**
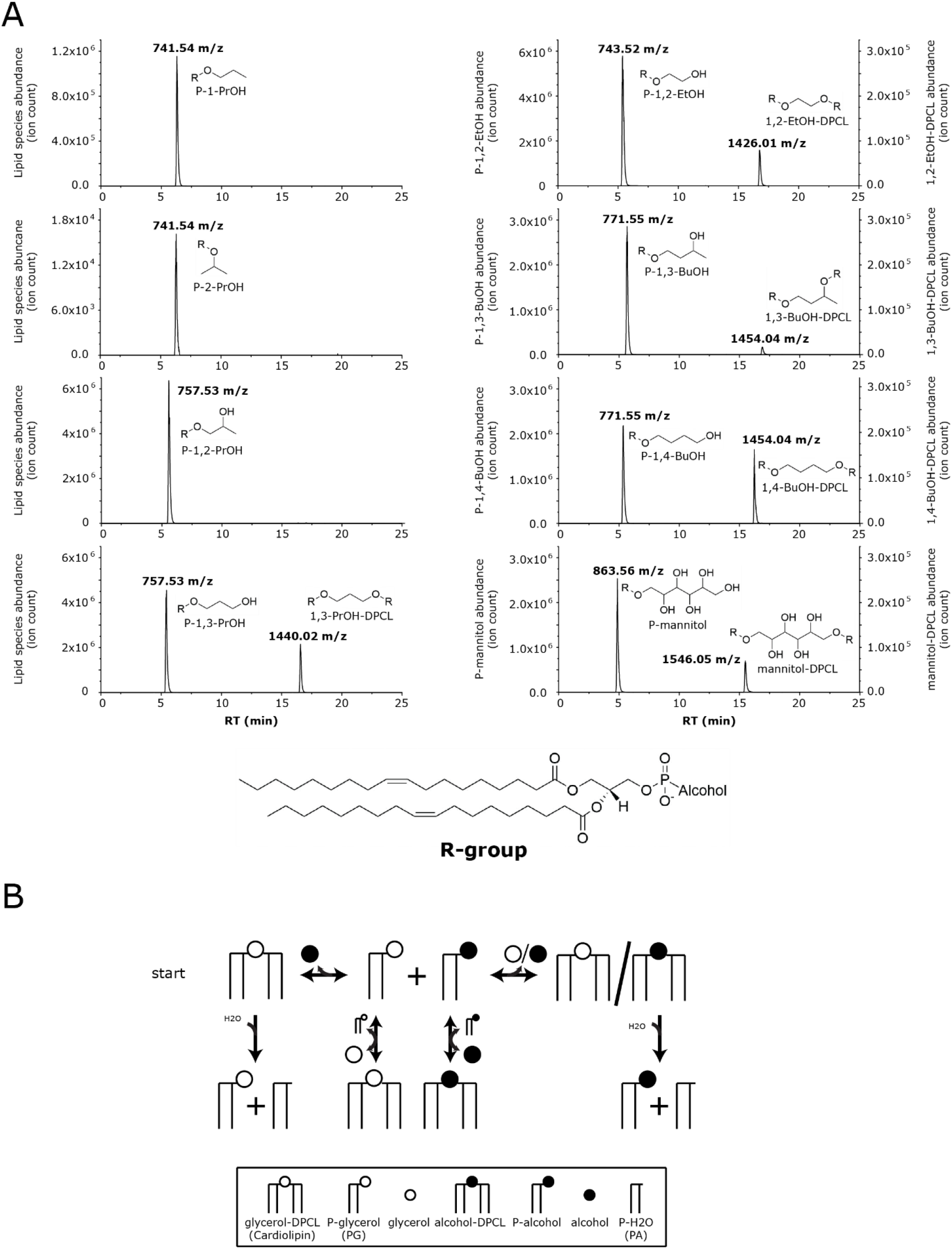
MhCls activity in the presence of various alcohols I. **(a)** Glycerol-like alcohols: 1-propanol, 2-propanol, 1,2-propanediol, 1,3-propanediol, 1,2-ethanediol, 1,3-butanediol, 1,4-butanediol and mannitol. **(b)** schematic representation of all possible reactions performed and products formed by MhCls, starting with cardiolipin and a substrate containing 2 primary hydroxyl groups (left top).

As in the presence of 1,2-propanol only trace amounts of 1,2-PrOH-DPCL could be detected, it seems that a second primary hydroxyl group is essential for the formation of a CL analogue. This was further tested with the substrates 1,2-ethanediol, 1,3-butanediol, and 1,4-butanediol. All three molecules functioned as a substrate for MhCls in the cardiolipin consuming reaction, which resulted in the production of the respective diester phospholipids, P-1,2-ethanediol (P-1,2-EtOH), P-1,3-butanediol (P-1,3-BuOH, and P-1,4-butanediol (P-1,4-BuOH). Moreover, for all these reaction conditions a DPCL equivalent could be detected as well. However, in the presence of 1,3-butanediol, instead of 1,4-butanediol, only 15% of butanediol-di-phosphate cardiolipin (BuOH-DPCL) could be produced, indicating that a primary hydroxyl group is preferred over a secondary hydroxyl group as a phosphatidyl-acceptor. This was further confirmed by the incorporation of mannitol (a six-carbon polyol), which resulted in the production of both phosphatidyl-mannitol (P-mannitol) and mannitol-di-phosphate cardiolipin (mannitol-DPCL), respectively. These experiments do not only show that the enzyme can utilize a wide variety of primary alcohols, but also exemplify its versatility. By starting with an isomerically pure, symmetric, cardiolipin and a substrate containing two primary hydroxyl groups, up to four additional lipid species can be synthesized (Figure 5B). The number of species can be further increased by using an isomerically pure, asymmetric cardiolipin instead, in which also variations in the acyl chain configuration contribute.

Next, we introduced molecules similar to glycerol, but with different bulky side-chains at the C-2 atom (Figure 6A). The presence of two methyl groups or a phenyl group at the C-2 position of the propanediol did not prevent these compounds from being used as a substrate, which resulted in the production of phosphatidyl-2,2-dimethyl-1,3-propanediol (P-2,2-Me-1,3-PrOH and phosphatidyl-2-phenyl-1,3-propanediol (P-2-Phe-1,3-PrOH). However, only trace amounts of the respective cardiolipin analogues could be formed, which indicates that these phosphatidyl-alcohol lipids do not function as suitable phosphatidyl-acceptors, possibly because of the steric hindrance of the substituents on the lipid headgroup.

**Figure 6:**
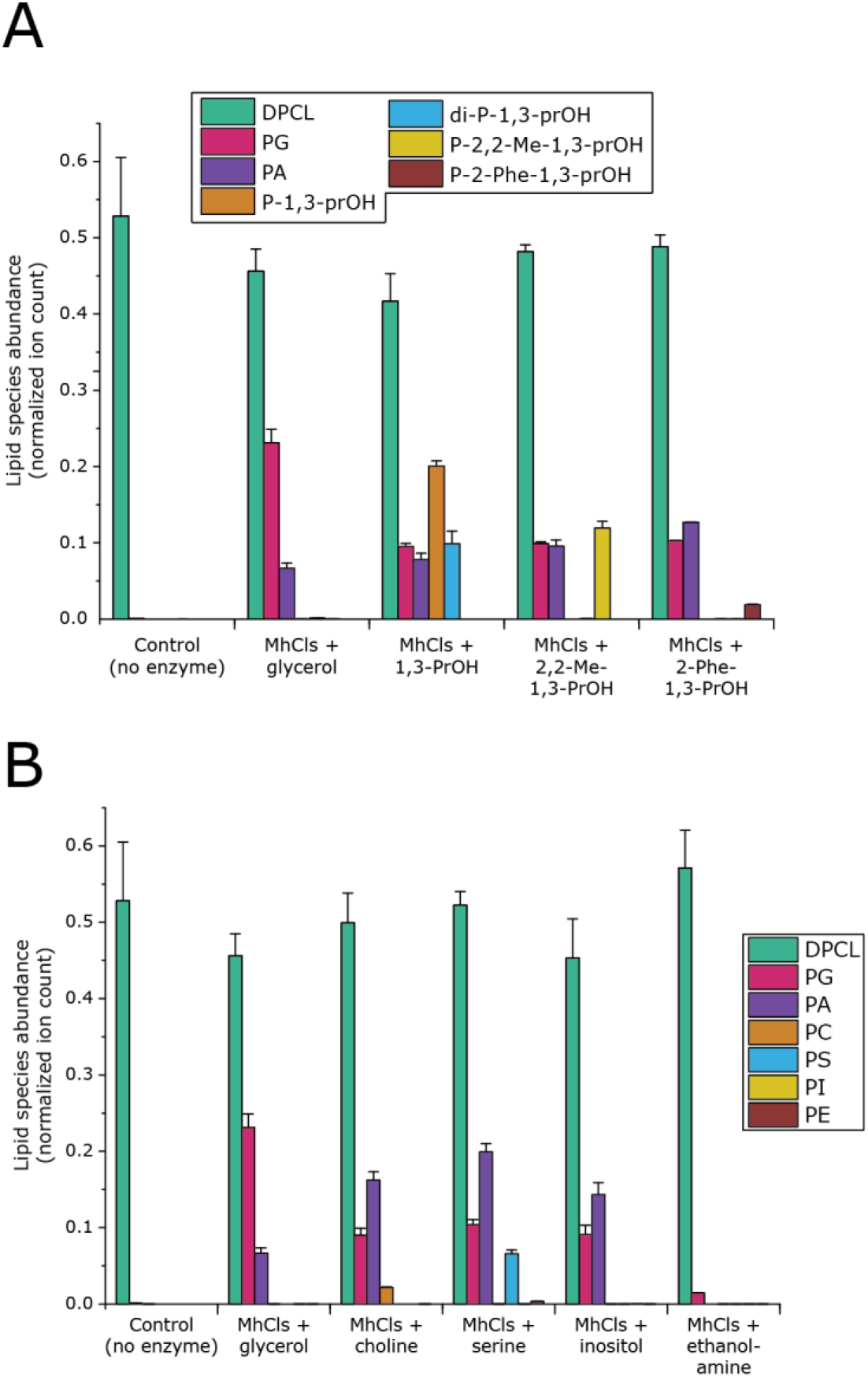
MhCls activity in the presence of various alcohols II. **(a)** 1,3-propanediol and its derivatives: glycerol (1,2,3-propanetriol), 2,2-dimethyl-1,3-propanediol and 2-phenyl-1,3-propanediol. **(b)** Common polar lipid headgroup alcohols: choline, serine, inositol, ethanolamine. Lipid species were analyzed by LC-MS, normalized for the internal standard DDM, and plotted.

Furthermore, a series of primary alcohols of biological relevance were tested. Herein, choline, L-serine, and ethanolamine were included in the reaction. In the case of choline and L-serine, the phospholipid species phosphatidylcholine (PC) and phosphatidylserine (PS) were formed together with PG and PA (Figure 6B). However, in the presence of ethanolamine, no phosphatidylethanolamine (PE) was formed. Notably, the production of PA and PG was largely abolished as well, suggesting that ethanolamine is an inhibitor of MhCls. An inhibiting effect was also observed in the presence of 3-amino-propanol (Figure S7A). Likewise, MhCls is unable to synthesize cardiolipin (analogues) or PA from a reaction mixture of PG in the presence of either 3-amino-propanol or ethanolamine, suggesting that primary amines inhibit the enzyme (Figure S7B). Finally, the sugar inositol was also tested even though this molecule does not contain any primary hydroxyl groups, and as expected no phosphatidylinositol (PI) was found.

### Glycocardiolipin formation

Since archaea also contain glycosyl-mono-phosphate cardiolipin (glyco-MPCL), the question arises if MhCls could also catalyze their synthesis (19). These molecules basically consist of a glycolipid ester-bonded to a phospholipid, thus containing only one instead of two phosphate moieties and a sugar headgroup instead of a glycerol (Figure 7A). To examine the ability of MhCls to make a glycol-MPCL, monogalactosyldiacylglycerol (MGDG), a glycolipid species present in the thylakoid membrane of higher plant chloroplasts, was tested as a possible substrate together with PG. Noteworthy, the MGDG used is a mixture of natural lipids with different fatty acid compositions. For simplicity we focused on the most abundant MGDG species (65-70%) with the acyl-chain configuration: 16:3-18:3 (*m/z* 745.49 [M-H]^−^). To promote glyco-MPCL formation over DPCL, MGDG was added in a two-fold excess compared to PG. However, since MGDG is a non-bilayer forming lipid, an excess of PC was also added to ensure bilayer formation for enzyme reconstitution, which resulted in a molar PG:MGDG:PC ratio of 1:2:17. In the presence of MhCls, the expected products, di-phosphate-cardiolipin (DPCL) and PA were formed in substantial amounts with concomitant utilization of PG (Figure. 7B). Furthermore, small amounts of another compound (*m/z* 1427.98 [M-H]^−^, mass error: 0.91 ppm) could be detected, corresponding to the glyco-MPCL species monogalactosyl-mono-phosphate cardiolipin (1Gal-MPCL) (Figure 7B). Although the signal is low, it is clearly detectable and elutes during the expected retention time range. MGDG elutes earlier from the column compared to PG, and therefore it is expected that the retention time of 1Gal-MPCL is also shorter than that of DPCL (Figure S8). Moreover, this mass is not present in the control condition without enzyme (Figure 7B), or without MGDG (data not shown), confirming that this molecule can only be formed by MhCls in the presence of MGDG.

**Figure 7:**
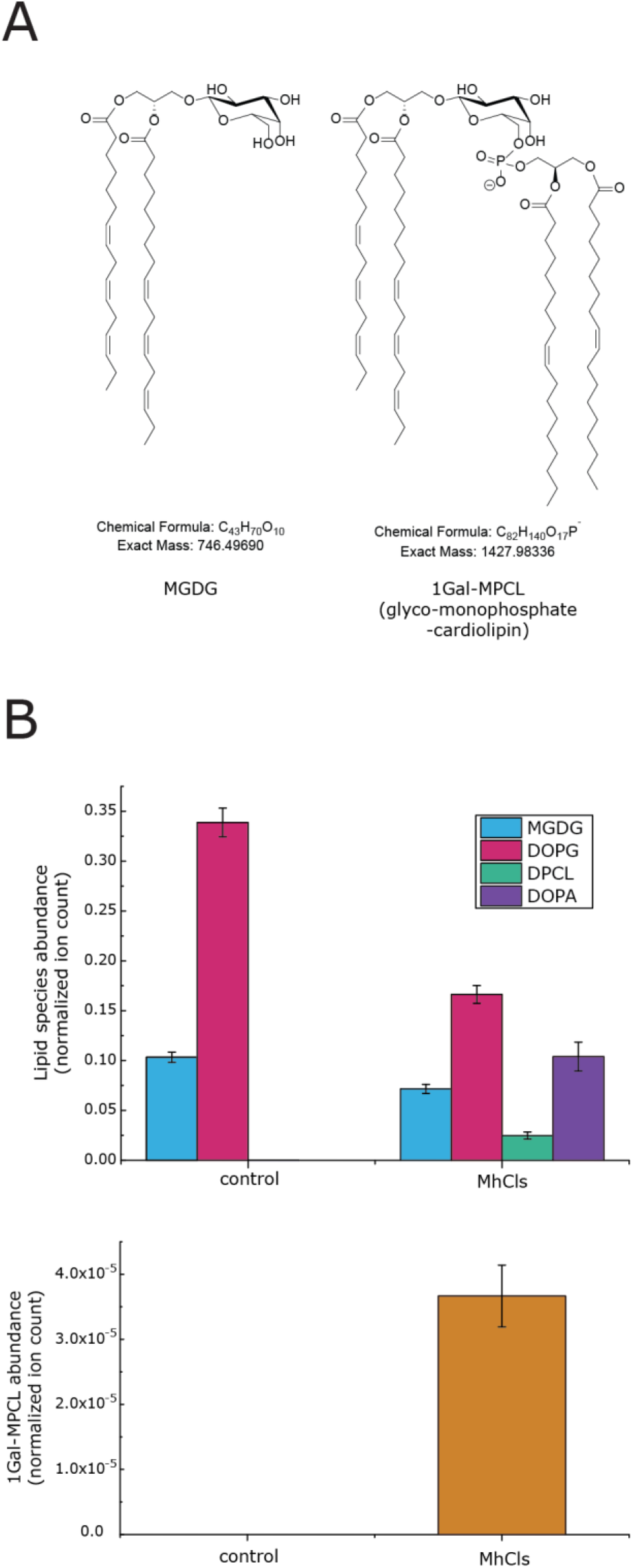
MhCls-dependent glycocardiolipin formation. **(a)** Structures of glycolipid MGDG and glycosyl-mono-phosphate cardiolipin 1Gal-MPCL. **(b)** MhCls mediated formation of DPCL, DOPA and 1Gal-MPCL. Lipid species were analyzed by LC-MS, normalized for the internal standard DDM, and plotted.

## Discussion

Cardiolipin (DPCL) is a lipid species found in the membranes of all domains of life. In contrast to eukaryotes and bacteria, the enzymes responsible for cardiolipin biosynthesis have not been studied in Archaea. Here, we identified the cardiolipin synthase from *Methanospirillum hungatei* (MhCls), which is a member of the phospholipase D superfamily (29), and characterized its function. Archaeal phospholipids consist of isoprenoid tails that are ether-linked to a glycerol-1-phosphate backbone. As a result, their chemical composition and chirality differ from the bacterial and eukaryotic lipids made of fatty acid-tails that are coupled to a glycerol-3-phosphate backbone via an ester-bond. Nevertheless, MhCls indiscriminately uses AG or PG to generate aDPCL and DPCL, respectively. This promiscuous feature of the enzyme enables it to simultaneous utilize PG and AG, resulting in the production of a novel hybrid-cardiolipin (hDPCL) species that contains one archaetidyl- and one phosphatidyl-moiety bridged by a glycerol headgroup. So far, such a molecule has not been observed in natural membranes, which can be attributed to an evolutionary event in which our last universal common ancestor (LUCA) evolved into the domains of Archaea and Bacteria. This event is also known as the “lipid divide” and distinguishes both domains with respect to the chemical composition and chirality of their membrane phospholipids (30, 31). The ability of MhCls to synthesize aDPCL, DPCL and hDPCL implies that critical substrate recognition only involves the polar headgroup of AG and PG (32, 33).

During the *in vitro* formation of cardiolipin, substantial levels of archaetidic acid (AA) and phosphatidic acid (PA) were noted, that emerged from the hydrolytic degradation of aDPCL and DPCL, respectively. As a consequence, cardiolipin synthase may act in lipid remodeling. For instance, the PA produced through the hydrolysis of DPCL, can be re-utilized by the enzyme CDP-diacylglycerol synthase (CdsA), for the formation of other phospholipid species (34, 35). The same accounts for the produced AA, which can be recycled back into the lipid biosynthesis route by CDP-archaeol synthase CarS (36). However, *in vitro* in the presence of only MhCls, PA cannot be re-utilized and accumulates in time, eventually depleting the PG-CL pool.

Our data show that MhCls catalyzes a glycerol-dependent dynamic equilibrium between its product DPCL, and the substrate PG. However, the enzyme exhibits a remarkable substrate promiscuity towards the lipid head group. Besides glycerol (and H_2_O), MhCls can incorporate various other substrates in the cardiolipin-utilizing reaction, yielding a wide variety of phosphatidyl-containing lipid species. Various primary alcohols can be attached to a phosphatidyl-group in the DPCL utilizing reaction, which results in the formation of PG and the specific phosphatidyl-alcohol. This is illustrated by the ability of MhCls to incorporate ethanediol, a two-carbon-diol, as well as the six-carbon-polyol mannitol, showing the enzyme its flexibility towards the length of the carbon-chain. Moreover, 1,3-propanediol derivatives with varying bulky side-groups at the second carbon can serve as phospholipid head group as well. Besides that, compounds with more biological relevance were tested, in which PC and PS could be formed during the conversion of DPCL in the presence of choline and serine, respectively. However, substrates that contain a primary amine (e.g. ethanolamine), act as inhibitors of the activity of the enzyme both in the cardiolipin biosynthetic and hydrolytic reaction. In addition, some phosphatidyl-alcohol species that are synthesized from a substrate that contain a second primary hydroxyl-group, can be further converted into a cardiolipin-analogue with the alcohol as bridging head group, thereby forming atypical and non-natural cardiolipins. The versatility of MhCls is further exemplified by the production of the glycosyl-mono-phosphate cardiolipin 1Gal-MPCL from the glycolipid substrate MGDG and the phospholipid substrate PG. This molecule was only detected in small amounts, indicating that MGDG (originating from plants) is a poor substrate, but the data supports the notion that this enzyme can produce a glycocardiolipin and that their synthesis might not involve a separate class of enzymes. Thus in halophiles, the identified glycosyl-mono-phosphate cardiolipins likely arise from a reaction that involves archaetidylglycerol and the respective glycolipid precursor (20–22).

Our *in vitro* assays show that MhCls can perform a wide variety of catalytic reactions. They are all based on the reversible transfer of a primary alcohol to a phosphatidyl-group. The implications of this promiscuity for the biological function of MhCls is as yet unclear. So far, no cardiolipins have been reported in the *M. hungatei* lipidome (37, 38), but those lipidomics studies were performed under growth conditions where the m*hCls* gene is barely expressed (39). This is not an uncommon finding as in many bacteria and, insofar studied, archaea the cardiolipin synthases are predominantly expressed during specific conditions (e.g. osmotic shock, specific growth phase, etc.), yielding different levels of cardiolipin (15–18). In this respect, the observed reversibility of the phosphatidyl-transfer makes the abundance of CL flexible, which could aid in the environmental response. Moreover, the adaptive ability of the membrane might be diversified with the promiscuous behavior of MhCls, illustrated by the ability of the enzyme to accept a wide variety of primary alcohols and lipids, which could be a general feature for cardiolipin synthases (25, 40–42).

Finally, the promiscuity of MhCls could be utilized for bioengineering purposes. As an example, the ability of this enzyme to incorporate a wide variety of non-natural polar head groups into phospholipid species may lead to new bio-catalytic applications for the synthesis of unique phospholipid species. Furthermore, MhCls potentially could be used for the bottom-up construction of a synthetic cellular membrane, in which this enzyme could diversify the phospholipid head group composition of an expanding phospholipid bilayer (34).

## Materials and methods

### Bioinformatic identification of MhCls

Using *E. coli* K12; MG1655 ClsA (EcClsA: NP_415765.1), ClsB (EcClsB: WP_187790083), or ClsC (EcClsC: WP_188006884.1) as query sequences, BLAST homology searches to the domain of Archaea were performed with the following result: EcClsA: query coverage 75-98%, and sequence identity 23-31%; EcClsB: query coverage 76-88%, and sequence identity 25-33%; EcClsC: query coverage 69-91%, and sequence identity 21-28%. The blast results were further analyzed using MEGA X and filtered for sequences that contain at least two HKD domains. Next, sequences were aligned using the MUSCLE algorithm (default settings) and a phylogenetic tree was estimated using the LG+G model (43). Subsequently, a putative archaeal cardiolipin synthase (MhCls: WP_011448254) from *Methanospirillum hungatei* JF-1 was selected. The consensus sequence logo was created with the program WebLogo (https://weblogo.berkeley.edu/logo.cgi), for which the group of methanogens was selected together with a group of bacterial EcClsA homologs (see supporting information). This resulted in a group of 90 species (57 bacteria and 33 archaea), from which one archaeal sequence, containing many additional amino acids within the second hydrophobic domain, was removed.

### Bacterial strains and cloning procedures

An *E. coli* codon-optimized synthetic gene of *M. hungatei* Cls (MhCls) was ordered (GeneArt, Thermo Fisher scientific) and used as a template for the amplification of MhCls, during which a N-terminal 6His-tag was added. The 6His-MhCls fragment was cloned into pRSF-Duet using the NcoI and SacI restriction enzymes and T4 DNA ligase resulting in pRSF-6His-MhCls. *E. coli* DH5α (Invitrogen) was used as a host for Cloning procedures. All primers and plasmids used in the present study are listed in Tables 1 and 2. *E. coli* Lemo21 (DE3) was used as the overexpression strain for MhCls. All strains were grown under aerobic conditions at 37°C in LB medium supplemented with the required antibiotics, kanamycin (50 μg/ml) and chloramphenicol (34 μg/ml).

**Table 1.**
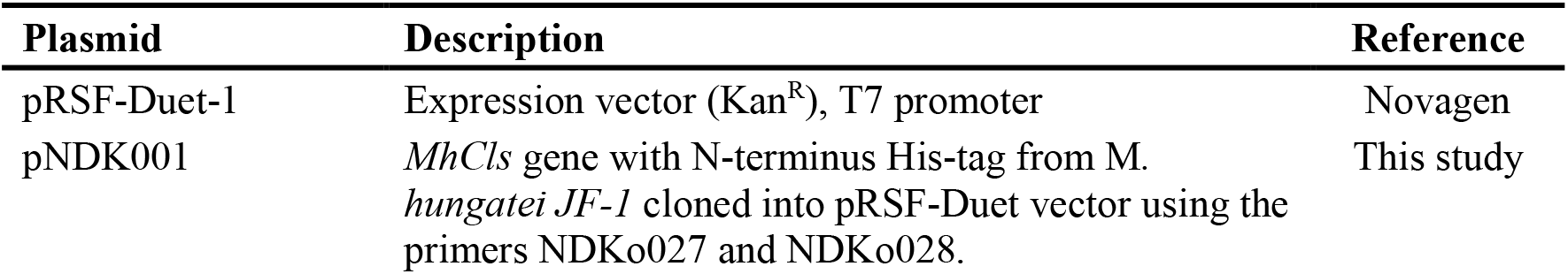
Cloning and expression vectors used in this study.

| Plasmid | Description | Reference |
| --- | --- | --- |
| pRSF-Duet-1 | Expression vector (Kan<sup>R</sup>), T7 promoter | Novagen |
| pNDK001 | *MhCls* gene with N-terminus His-tag from *M. hungatei JF-1* cloned into pRSF-Duet vector using the primers NDKo027 and NDKo028. | This study |

**Table 2.**
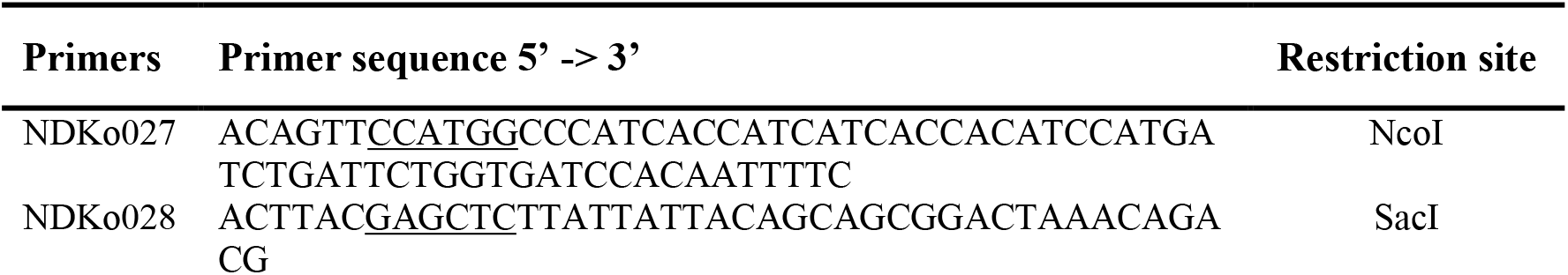
Oligonucleotide primers used in this study.

### Expression and purification of MhCls

MhCls was overexpressed in *E. coli* Lemo21 (DE3) strain in the presence of 250 μM rhamnose and induced with 0.5 mM isopropyl β-D-1-thiogalactopyranoside (IPTG). After 2.5 hours of induction, cytoplasmic and membrane fractions were separated as described (44). The total membranes were resuspended in buffer A (50 mM Tris/HCl, pH 8.0, 100 mM KCl and 15% glycerol) after which they could be stored at −80°C. For further purification, 0.5 mg/ml of membranes were solubilized in 2% n-dodecyl-β-D-maltopyranoside (DDM) detergent for 1 hr. at 4°C. The material was subjected to a centrifugation (17,000 x g) step for 15 min at 4°C to remove insolubilized material and the supernatant was incubated with Ni-NTA agarose beads (Qiagen, cat: 30230) for 2 hrs. at 4°C. The Ni-NTA beads were washed 10 times with 6 column volumes (CV) of buffer B (50 mM Tris/HCl, pH 8.0, 100 mM KCl, 15% glycerol and 0.05% DDM) supplemented with 10 mM imidazole, and the protein was eluted three times with 0.5 CV of buffer B supplemented with 300 mM imidazole. To remove the imidazole and glycerol, the purified protein was passed over a Zeba™ Spin Desalting column 40K; 0.5 ml (Thermo scientific), and eluted in buffer C (50 mM MES pH 7.0, 100 mM KCl and 0.05% DDM). Purity of the eluted protein was assessed on 15% SDS/PAGE stained with Coomassie Brilliant Blue and the protein concentration was determined by measuring the absorbance at 280 nm and calculating the molar concentration using the calculated extinction coefficient. Extinction coefficients were obtained from the ProtParam tool from the ExPASy website (https://web.expasy.org/protparam/).

### Liposomes preparation

Chloroform stocks of the lipid species DOPG, POPG, DOPA, POPA, DOPE, DOPC, DPCL 18:1/18:1/18:1/18:1 and MGDG were purchased from Avanti (Avanti Biochemicals, Birmingham, AL). The chemical synthesis of AG was performed in house and is described in detail in the supporting information. For liposomes with a heterogeneous lipid mixture, the required amount of lipid chloroform stocks were mixed together in the stated molar ratio. Next the lipid solution was dried under a nitrogen gas stream for multiple hours, after which the dry lipid film was resuspended in a 50 mM 2-(N-morpholino)ethanesulfonic acid (MES) buffer, pH 7.0 yielding a translucent suspension. For formation of liposomes, a probe sonicator was employed (30 seconds cycle time with a 50% duty cycle for 10-20 cycles) until the suspension became transparent.

### *In vitro* assays for phospholipid production

All *in vitro* reactions were performed in 100 μl of buffer D containing a final concentration of 50 mM MES pH 7.0 and 100 mM KCl in the presence of 1 μM MhCls. The activity of MhCls with archaeal substrate was assayed in the presence of 250 μM AG (*sn*1-*sn*1 and *sn*1-*sn*3; ratio 1:1) and 0.4 mM DDM. The glycerol-dependent dynamic equilibrium of MhCls was assayed in the presence of 1.8 mM DDM, and either 1 mM DOPG, 0.5 mM DPCL, or 0.5 mM POPG together with 0.25 mM DPCL. The activity of MhCls with AG and PG (molar ratio 1:1) was assayed with 250 μM of each lipid substrate in the presence of 0.8 mM DDM. The promiscuity toward primary-hydroxyl-containing compounds was assayed with 0.5 mM DPCL or 1 mM DOPG, 100 mM primary-hydroxyl-containing substrate and 1.8 mM DDM. Formation of glycosyl-mono-phosphate cardiolipin was performed in the presence of 1 μM MhCls, 0.5 mM lipid (PG:MGDG:PC, molar ratio 1:2:17) and 0.8 mM DDM. All reactions were incubated overnight at 37°C unless stated differently. Lipids were extracted from the reaction mixtures two times with 0.3 ml of 1-butanol, and evaporated under a stream of nitrogen gas and resuspended in 50 μl of methanol for LC-MS analysis.

### LC–MS analysis of lipids

Samples from the *in vitro* reactions were analyzed using an Accela1250 high-performance liquid chromatography (HPLC) system coupled with an Heated electrospray ionization mass spectrometry (HESI–MS) Orbitrap Exactive (Thermo Fisher Scientific). A sample of 5 μl was injected into an ACQUITY UPLC® CSH™ C18 1.7 μm Column, 2.1×150 mm (Waters Chromatography Ireland Ltd) operating at 55°C with a flow rate of 300 μl/min. Separation of the compounds was achieved by a changing gradient of eluent A (5 mM ammonium formate in water/acetonitrile 40:60, v/v), and eluent B (5 mM ammonium formate in acetonitrile/1-butanol, 10:90, v/v). The following linear gradient was applied: 45% eluent B for 2.5 minutes; a gradient from 45% to 90% eluent B over 19.5 minutes; holding for 3 minutes; returning to 45% eluent B in 0.5 minutes; and holding for 8 minutes. The column effluent was injected directly into the Exactive ESI-MS Orbitrap operating in negative ion mode. Voltage parameters of 3 kV (spray), −75 V (capillary), −190 V (tube lens) and −46 V (Skimmer voltage) were used. Capillary temperature of 300°C, sheath gas flow 60, and auxiliary gas flow of 5 was maintained during the analysis.

Spectral data constituting total ion counts were analyzed using the Thermo Scientific XCalibur processing software by applying the Genesis algorithm based automated peak area detection and integration. The total ion counts of the extracted lipid products (Table 3) were normalized for DDM (*m/z* 509.3 [M-H]^−^) and plotted on the y-axis as normalized ion count in a bar graph.

**Table 3.** detected lipid species with LC-MS

| Lipid species | *m/z*<br>[M-H]<sup>-</sup> |
| --- | --- |
| archaetidylglycerol (AG) | 805.66 |
| archaetidic acid (AA) | 731.63 |
| archaetidic di-phosphate cardiolipin (aDPCL) | 1520.30 |
| di-oleoyl phosphatidic acid (DOPA), phosphatidic acid (PA) | 699.49 |
| di-oleoyl phosphatidylglycerol (DOPG), phosphatidylglycerol (PG) | 773.53 |
| palmitoyl-oleoyl phosphatidic acid (POPA) | 673.48 |
| palmitoyl-oleoyl phosphatidylglycerol (POPG) | 747.52 |
| di-phosphate-cardiolipin (DPCL), DPCL18:1/18:1/18:1/18:1 | 1456.03 |
| DPCL16:0/18:1/16:0/18:1 | 1403.99 |
| DPCL18:1/18:1/16:0/18:1 | 1430.01 |
| hybrid di-phosphate cardiolipin (hDPCL) | 1488.16 |
| phosphatidyl-1-propanol (P-1-PrOH), and phosphatidyl-2-propanol (P-2-PrOH) | 741.54 |
| phosphatidyl-1,2-propanediol (P-1,2-PrOH), and phosphatidyl-1,3-propanediol (P-1,3-PrOH) | 757.53 |
| 1,2-propanediol-di-phosphate cardiolipin (1,2-PrOH-DPCL), and 1,3-propanediol-di-phosphate cardiolipin (1,3-PrOH-DPCL) | 1440.02 |
| phosphatidyl-1,2-ethanol (P-1,2-EtOH) | 743.52 |
| 1,2-ethanol-di-phosphate cardiolipin (1,2-EtOH-DPCL) | 1426.01 |
| phosphatidyl-1,3-butanediol (P-1,3-BuOH), and phosphatidyl-1,4-butanediol (P-1,4-BuOH) | 771.55 |
| 1,3-butanediol-di-phosphate cardiolipin (1,3-BuOH-DPCL), and 1,4-butanediol-di-phosphate cardiolipin (1,4-BuOH-DPCL) | 1454.04 |
| phosphatidyl-mannitol (P-mannitol) | 863.56 |
| mannitol-di-phosphate cardiolipin (mannitol-DPCL) | 1546.05 |
| phosphatidyl-2,2-dimethyl-1,3-propanediol (P-2,2-Me-1,3-PrOH) | 786.56 |
| phosphatidyl-2-phenyl-1,3-propanediol (P-2-Phe-1,3-PrOH) | 833.56 |
| di-oleoyl phosphatidylethanolamine (DOPE) | 742.54 |
| phosphatidyl-aminopropanol (P-NH<sub>2</sub>-PrOH) | 756.55 |
| di-oleoyl phosphatidylcholine (DOPC) | 830.59 |
| di-oleoyl phosphatidylserine (DOPS) | 786.53 |
| di-oleoyl phosphatidylinositol (DOPI) | 861.55 |
| monogalactosyldiacylglycerol (MGDG) | 746.49 |
| monogalactosyl-di-phosphate cardiolipin (1Gal-MPCL) | 1427.98 |

## Data availability

All raw and processed data used for and described in this article is stored in the department of Molecular Microbiology at the University of Groningen.

## Author contributions

Conceptualization, M.E., N.A.W.K., and A.J.M.D.; Investigation: M.E., N.A.W.K., R.L.H.A., N.H.J.W.; Resources: A.J.M., A.J.M.D.; Writing ‒Original Draft: M.E., R.L.H.A.; Writing ‒Review & Editing: M.E., N.A.W.K., A.J.M., A.J.M.D.; Visualization: M.E., R.L.H.A.; Supervision, M.E., A.J.M., A.J.M.D.; Project Administration, M.E.; Funding Acquisition, A.J.M.D., A.J.M.;

## Funding and additional information

This work was supported and funded by the ‘BaSyC – Building a Synthetic Cell’ Gravitation grant (024.003.019) of the Netherlands Ministry of Education, Culture and Science (OCW) and the Netherlands Organization for Scientific Research (NWO), and by the Dutch NWO Building Blocks of Life programme (737.016.006).

## Abbreviations

AA: archaetidic acid
aDPCL: archaetidic di-phosphate cardiolipin
AG: archaetidylglycerol
CDP-DAG: cytidine diphosphate diacylglycerol
CL: cardiolipin
Cls: cardiolipin synthase
ClsA: cardiolipin synthase A
ClsB: cardiolipin synthase B
ClsC: cardiolipin synthase C
DDM: n-dodecyl-β-d-maltoside
DOPA: di-oleoyl phosphatidic acid
DOPC: di-oleoyl phosphatidylcholine
DOPE: di-oleoyl phosphatidylethanolamine
DOPG: di-oleoyl phosphatidylglycerol
DOPI: di-oleoyl phosphatidylinositol
DOPS: di-oleoyl phosphatidylserine
DPCL: di-phosphate-cardiolipin
glyco-MPCL: glycosyl-mono-phosphate cardiolipin
hDPCL: hybrid di-phosphate cardiolipin
LUCA: last universal common ancestor
mannitol-DPCL: mannitol-di-phosphate cardiolipin
MGDG: monogalactosyldiacylglycerol
MhCls: *Methanospirillum hungatei* cardiolipin synthase
Ni-NTA: Nickel-nitrilotriacetic Acid
PA: phosphatidic acid
PG: phosphatidylglycerol
POPA: palmitoyl-oleoyl phosphatidic acid
POPG: palmitoyl-oleoyl phosphatidylglycerol
P-mannitol: phosphatidyl-mannitol
P-NH2-PrOH: phosphatidyl-aminopropanol
P-1-PrOH: phosphatidyl-1-propanol
P-1,2-EtOH: phosphatidyl-1,2-ethanol
P-1,2-PrOH: phosphatidyl-1,2-propandiol
P-1,3-BuOH: phosphatidyl-1,3-butanediol
P-1,3-PrOH: phosphatidyl-1,3-propandiol
P-1,4-BuOH: phosphatidyl-1,4-butanediol
P-2-Phe-1,3-PrOH: phosphatidyl-2-phenyl-1,3-propanediol
P-2-PrOH: phosphatidyl-2-propanol
P-2,2-Me-1,3-PrOH: phosphatidyl-2,2-dimethyl-1,3-propanediol
S-DGD-5-PA: S-diphytanylglycerol diether-5-phosphatidic acid
S-2Glyco-aMPCL: S-di-glycosyl-archaeal mono-phosphate cardiolipin
S-2Glyco-DGD: S-di-glycosyl diphytanylglycerol diether
S-3Glyco-aMPCL: S-tri-glycosyl-archaeal mono-phosphate cardiolipin
S-GL-2: S-glycosylcardiolipin-2
S-TGD-1-PA: S-tri-glycosyl-diether-1-phosphatidic acid
1Gal-MPCL: monogalactosyl-di-phosphate cardiolipin
1,2-EtOH-DPCL: 1,2-ethanol-di-phosphate cardiolipin
1,2-PrOH-DPCL: 1,2-propanediol-di-phosphate cardiolipin
1,3-BuOH-DPCL: 1,3-butanediol-di-phosphate cardiolipin
1,3-PrOH-DPCL: 1,3-propanediol-di-phosphate cardiolipin
1,4-BuOH-DPCL: 1,4-butanediol-di-phosphate cardiolipin

